# Congress of multiple dimers is needed for cross-phosphorylation of IRE1α and its RNase activity

**DOI:** 10.1101/2023.12.18.572154

**Authors:** Andrea Orsi, Roberto Sitia, Eelco van Anken, Milena Vitale, Anush Bakunts

## Abstract

The unfolded protein response can switch from a pro-survival to a maladaptive, pro-apoptotic mode. During endoplasmic reticulum (ER) stress, IRE1α sensors dimerize, are phosphorylated and activate XBP1 splicing, increasing folding capacity in the ER protein factory. The steps that turn the IRE1α endonuclease activity against endogenous mRNAs during maladaptive ER stress are still unknown. Here we show that although necessary, IRE1α dimerization is not sufficient to trigger phosphorylation. Random and/or guided collisions amongst IRE1α dimers are needed to elicit cross-phosphorylation and endonuclease activities. Thus, reaching a critical concentration of IRE1α dimers in the ER membrane is a key event. Formation of stable IRE1α clusters is not necessary for RNase activity. However, clustering could modulate the potency of the response promoting interactions between dimers and decreasing the accessibility of phosphorylated IRE1α to phosphatases. The stepwise activation of IRE1α molecules and their low concentration at steady state prevent excessive responses, unleashing full-blown IRE1 activity only upon intense stress conditions.

## Introduction

Reliability of signal transduction is crucial for cell function and survival. The vast majority of secretory proteins fold and assemble in the endoplasmic reticulum (ER) under the assistance of resident chaperones and enzymes. Folding intermediates are retained in the early secretory compartment until they reach their native conformation (Anelli and Sitia, 2008). Proteins which fail to do so are cleared from the ER, most often through ER-associated degradation (ERAD), which entails dislocation to the cytosol for proteasome degradation (Ellgaard and Helenius, 2003, Sitia and Braakman, 2003; Krshnan et al., 2022). When the load of clients overwhelms the folding capacity of the protein factory, ER stress ensues, which in turn activates three adaptive pathways (Perk, ATF6, IRE1α) collectively referred to as the unfolded protein response (UPR) (Walter and Ron, 2011). The most conserved branch of the UPR is the one orchestrated by IRE1α. Upon ER stress, IRE1α is phosphorylated, oligomerizes and acquires endonuclease activity, yielding spliced XBP1 mRNA (XBP1s). The resulting sXBP1 protein is a transcription factor that drives the expression of ER resident chaperones, enzymes and ERAD components (Walter and Ron, 2011). In such capacity IRE1α plays a beneficial role as it is committed to re-establish ER homeostasis. However, under certain conditions, such as unresolved ER stress, IRE1α cleaves other RNAs in a process named regulated IRE1-dependent decay (RIDD, Hollien et al., 2009), which can cause apoptosis. Thus, a strict control of IRE1α activity is essential as inadequate regulation of the enzyme (both overactivation and underactivation) may lead to the premature death of otherwise healthy cells, or the survival of cells that instead ought to be eliminated. Its central role in cell life-death decisions is key in pathological processes such as cancer and diabetes, making IRE1α a promising therapeutic target (Raymundo et al., 2020, Morita et al., 2017).

Several reports have shown that upon intense stress Ire1p forms clusters or foci, which have been proposed to help recruitment of HAC1 mRNA (yeast homologous of XBP1) or have a role in the acquisition of IRE1α RNase activity (Aragón et al., 2009, van Anken et al., 2014). However, much remains to be understood on the chain of events that lead to and limit IRE1α activation. In this study, we analyzed a panel of mutants to dissect the stepwise role of dimerization, oligomerization, nucleotide binding, phosphorylation and RNase activity during progression of ER stress. We found that, surprisingly, IRE1α phosphorylation does not occur within isolated dimers, but only in trans upon collisions of dimers and/or formation of higher order oligomers. Owing to the low abundance of IRE1α, isolated dimers will bump into each other only occasionally – resulting in limited cross-phosphorylation. Thus, we infer that full-blown activation could be achieved only by formation of higher oligomers or by concentration of IRE1 dimers in specialized structures. The transition between dispersed, mildly activated IRE1 and fully active IRE1 in specialized foci may be crucial for controlling life/death cell decisions.

## Results and discussion

### IRE1α phosphorylation correlates in time and magnitude with its RNAse activity

We recently developed and validated a robust cell model, which allows evaluation of ER homeostatic readjustments in response a proteostatic insult, i.e. the overexpression of orphan secretory Ig-µ_s_ heavy chain (µ_s_, Bakunts et al., 2017; Vitale et al., 2019). Synthesis of exuberant levels of µ_S_ results in temporary shortage of free BiP, which leads to UPR activation. Using this model, we showed that reaching a new homeostatic equilibrium entails the transition from acute UPR signaling, when ER stress sensors are fully activated, to a chronic state characterized by an overall ER expansion (Bakunts et al., 2017; Vitale et al., 2019).

Here we exploited our model to follow IRE1α phosphorylation at different stages of a proteostatically-driven UPR. As previously described (Bakunts et al., 2017), at the early time-points corresponding to acute UPR a significant portion of IRE1α is phosphorylated and high levels of spliced XBP1 are generated (Fig 1A, B). Later, when a new homeostatic equilibrium is established, IRE1α phosphorylation subsides to levels close to those in steady-state and its RNAse activity decreases significantly (Fig 1A, B). However, when ER stress cannot possibly be resolved – i.e. because ERAD is blocked with kifunensine (kif) – the levels of IRE1α phosphorylation and XBP1 splicing remain high (Fig 1A, B) and cells eventually die. In this model, therefore, IRE1 phosphorylation and endonuclease activity parallel the intensity of stress.

**Fig 1.**
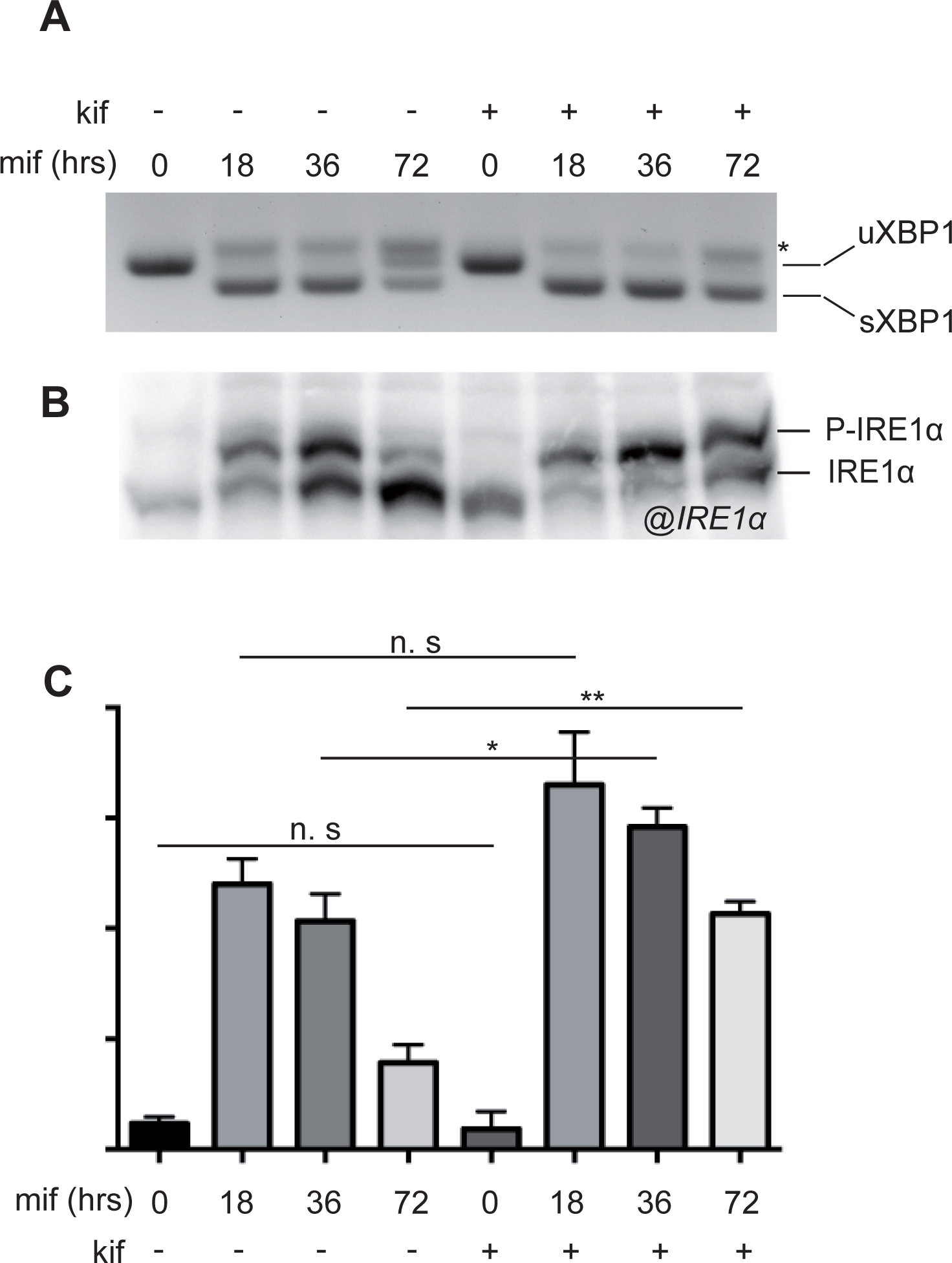
IRE1α phosphorylation level is proportional to its endonuclease activity. A. HeLa-µs cells were induced with 0.5 nM mifepristone (Mif) to induce Ig-µ chain synthesis and treated with or without 30 µM kifunensine (kif) for the indicated times, to induce adaptive or maladaptive UPR, respectively. XBP1 splicing was used as an indicator of IRE1α endonuclease activity. A hybrid product that is formed during the PCR reaction (Shang and Lehrman, 2004) is denoted by an asterisk. B. Aliquots from the lysates were resolved by Phos-tag gels and blots decorated with anti-IRE1α. See panel C for densitometric quantifications. C. The fraction of phosphorylated IRE1α was quantified by densitometry. Preventing ERAD by kifunensine addition increased the extent of IRE1α phosphorylation, particularly during the late phases of the response to Ig-µ chain synthesis. T-test compared portion of phosphorylated protein during Mif time-course in the presence and the absence of kif.

### A cellular model to investigate IRE1 activation

To dissect the molecular steps that lead to IRE1α activation, we generated CRISPR-knockout cells for IRE1α and reconstituted them with a panel of IRE1α mutants designed to pinpoint the roles and relationships between dimerization, oligomerization, phosphorylation, nucleotide-binding and full enzymatic activation (Fig 2A).

**Fig 2.**
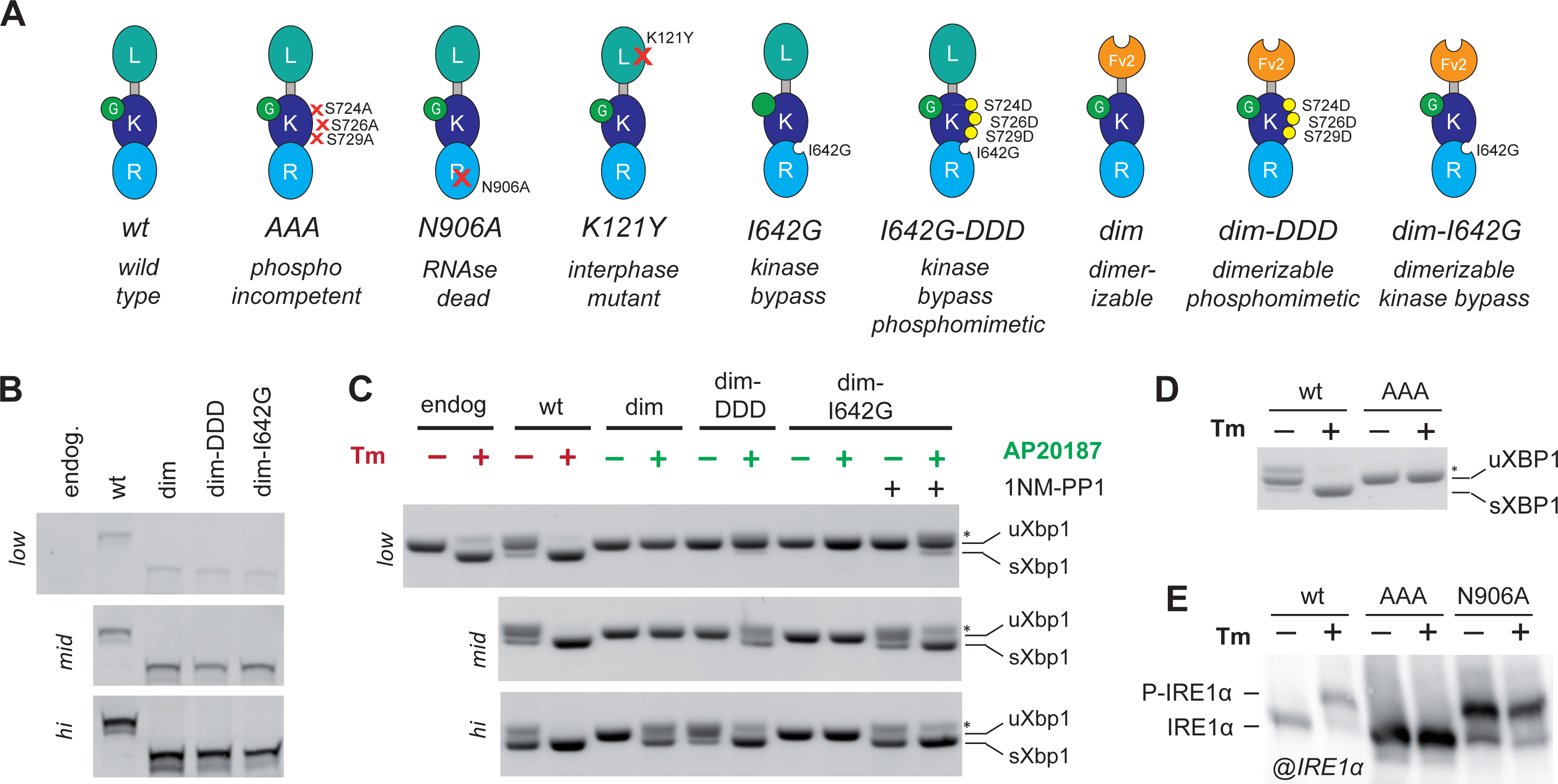
Dimerization is not sufficient for complete activation of IRE1α. A. The panel schematizes the IRE1a mutants used in the study. L: luminal domain; K: kinase domain; R: RNAse domain; Fv2E: artificially dimerizable domain. B. To visually compare the different (low, medium, high) levels of expression of the IRE1α mutants utilized, aliquots from the lysates were resolved electrophoretically and the blots decorated with anti-IRE1α. For higher exposure of “low” panel see Fig S1E. C. Comparison of the RNase activity of IRE1α mutants at different expression levels (low, medium, high) after treatment with or without tunicamycin or dimerizer drug (AP20187) and with or without 1NM-PP1 in case of Dim-I642G. Splicing of XBP1 was assessed as in Fig 1A. Clearly, the concentration of IRE1α is a key determinant in unleashing its endonuclease activity. D. Phosphorylation-deficient mutant (AAA) has no endonuclease activity even at high expression levels as assessed by the absence of sXBP1 after treatment with Tm. E. Aliquots from the lysates of cells expressing wt IRE1α or the AAA and N906A (RNAse dead) mutants at high levels treated with or without tunicamycin were resolved by Phos-tag gels and Blots were decorated with anti-IRE1α. While AAA cannot be phosphorylated in response to Tm, the N906A mutant is phosphorylated even in the absence of ER stress.

In order to compare the activity of the mutants it was essential to control their expression levels. To this end, we employed the TetON system and created a panel of inducible HeLa cell lines to express the different IRE1α mutants at three comparable levels: low, intermediate, and very high (Fig 2B, Fig S1A-E). In addition to that, all of our mutants are GFP-tagged to enable imaging studies.

First, we confirmed that full-length wild type IRE1α-GFP fully complements IRE1α KO cells. To monitor the phosphorylation status of IRE1α we employed both Phos-tag SDS-PAGE and specific anti-phosphoIRE1α antibodies. Upon treatment with tunicamycin (Tm), an inhibitor of N-glycosylation known to cause robust ER stress, IRE1α-GFP was efficiently phosphorylated and catalyzed XBP1 splicing, in a comparable way to endogenous IRE1α (Fig 2C, E).

The importance of IRE1α phosphorylation as a prerequisite for endonuclease activity is confirmed by analysis of a full-length phosphorylation-deficient IRE1α mutant (AAA) in which serines 724/726/729, located in the kinase activation loop of the protein, had been mutated to alanines. As expected, this mutant cannot be phosphorylated upon ER stress (Fig 2E). Different from previous reports (Prischi et al., 2014), in our hands the FL-AAA mutant was unable to splice XBP1 mRNA in response to ER stress. Endonuclease activity was not rescued even if the mutant was expressed at very high levels (Fig 2D, Fig S1A).

Conversely, IRE1α carrying N906A mutation, which abolish its RNAse activity (RNAse dead, RD) (Han et al., 2009) is phosphorylated even in the absence of ER stress (Fig 2E), confirming that phosphorylation precedes and can occur independently from the endonuclease activity of IRE1α.

### IRE1α dimers are not capable of auto-phosphorylation

Dimerization is known to be a prerequisite for IRE1α activation. In principle, one IRE1α dimer could be sufficient for a single XBP1 cleavage event (Korennykh et al., 2011). However, it is not clear if phosphorylation and the following steps require higher-order IRE1α oligomerization. Indeed, in several cellular models ranging from yeast to mammals, IRE1α was shown to form oligomers and big signaling clusters upon acute ER stress (Aragón et al., 2009, Korennykh A. and Walter P, 2012).

Tracking and controlling the transition between monomeric, dimeric and oligomeric forms of IRE1α in a living cell is quite challenging. To tackle this problem, we created a dimerizable IRE1α chimera (Dim-IRE1α) in which the luminal domain of IRE1α was replaced by a modified version of the FVBK domain (Fv2E). Two of such domains can be brought together artificially by addition of the divalent chemical AP20187, as it was done before for another ER stress sensor, Perk (Lin et al., 2009, Lu et al., 2004). Thanks to this chimeric construct, the transition between monomeric and dimeric IRE1α can be manipulated in a tightly controlled fashion, independently from ER stress and without concomitant activation of other UPR sensors. Moreover, the Dim-IRE1α mutant allows to assess the neat contribution of dimerization to phosphorylation and endonuclease activity, since this chimera lacks structural elements of the luminal domain which could mediate formation of stable IRE1α oligomers.

In the absence of the dimerizing agent, expression of Dim-IRE1α did not result in XBP1 splicing. Thus, appending the Fv2E domain does not mediate IRE1α activation *per se* (Fig 2C). Moreover, treatment with the dimerizing drug did not interfere with XBP1 splicing and IRE1α expression levels (Fig S2A, B) and the concentration of dimerizing drug was sufficient to saturate binding to the Fv2E domain (Fig S2C). Importantly, neither Dim-IRE1α phosphorylation nor XBP1 splicing was detected when the mutant was expressed at low and intermediate levels (Figs 2C and 3A). Notably, however, full-length wild type IRE1α was fully active when expressed at comparable levels. These findings suggested that the dimer itself is not capable of self-phosphorylation.

**Fig 3.**
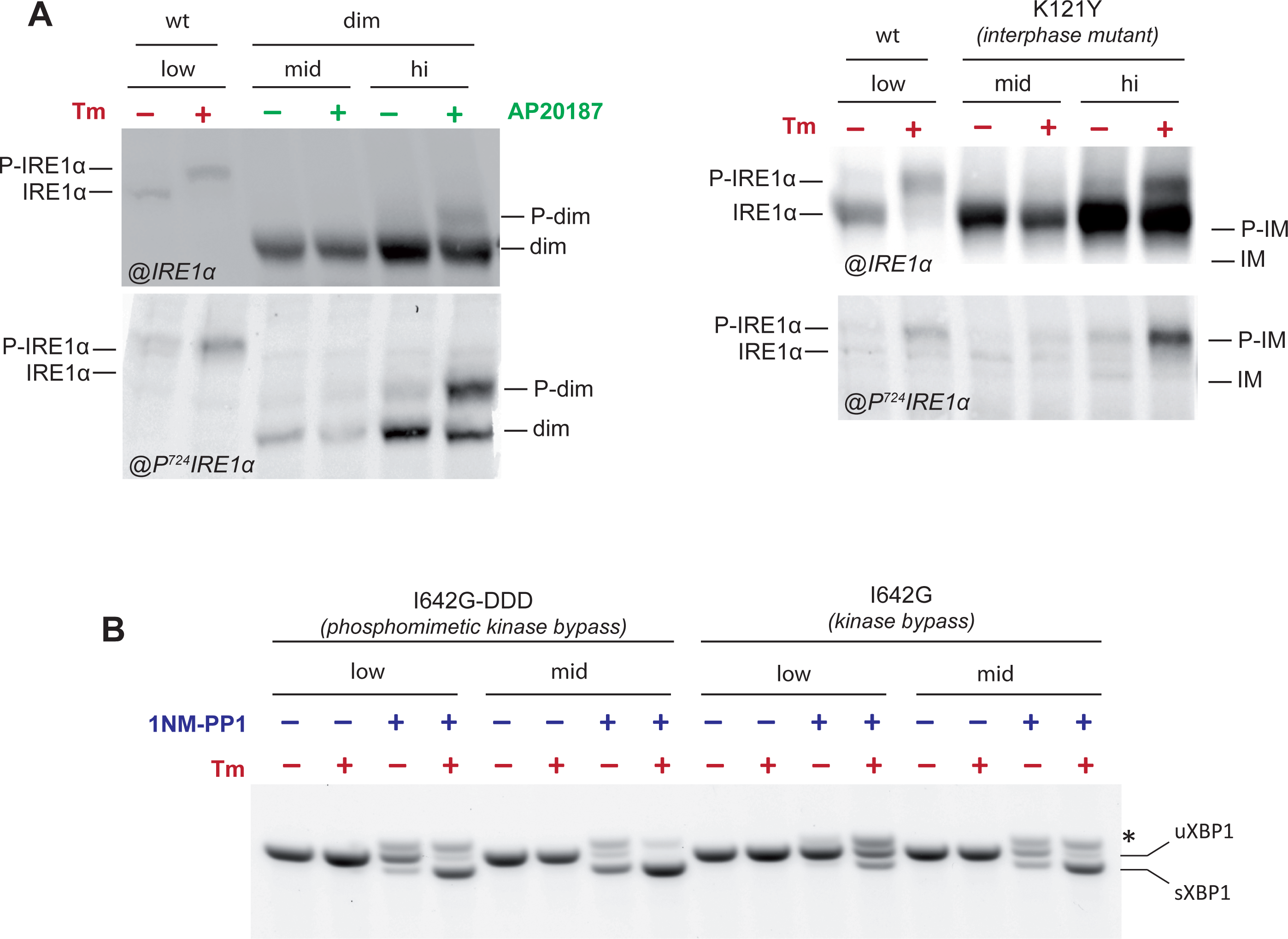
Higher order assemblies is necessary for IRE1α phosphorylation and activation. A. Aliquots from the lysates of cells treated with Tm or AP20187 as indicated were resolved on Phos-tag gels and immunoblotted with anti-IRE1α (upper panel) or anti-phosphoIRE1α (phospho S724, lower panel) antibodies. The first two lanes contain lysates from cells expressing wild type IRE1 at low levels. The remaining lanes show lysates from cells expressing dimerizable IRE1α (dim) at medium and high expression levels, as indicated. Both dimerizable IRE1α and full-length K121Y (interface mutant) are partially phosphorylated only at high expression levels. B. The panel show the endonuclease activity of the kinase bypass (I642G and I642G-DDD) nucleotide binding requiring IRE1α mutants, expressed at low and medium levels. The phosphomimetic IRE1α mutant I642G-DDD allows to evaluate the possible role of phosphorylation for stabilization of nucleotide-bound state. XBP1 splicing by both I642G mutants depends on nucleotide activation (compare for instance lanes 1-2 to 3-4), intracellular concentration (compare lanes 3 to 7 or 4 to 8) and phosphorylation (compare lanes 3 to 11 or 4 to 12). Splicing of XBP1 was assessed as in Fig 1A.

We also reasoned that the presence of the Fv2E domain might impair IRE1α kinase activity. To exclude this possibility, we generated a cell line expressing an interface mutant of IRE1α, which has an intact luminal domain but a single point mutation (K121Y) which impairs oligomerization propensity (Li et al., 2010; Sundaram et al., 2018). Again, this mutant was not active at close to endogenous expression levels (Fig S1C, F), indicating that reaching a higher-order oligomerization state is important to trigger phosphorylation. Importantly, due to the lack of the normal luminal domain, the Dim-IRE1α mutant cannot form foci. Also the K121Y mutation prevents IRE1α clustering in response to ER stress (Li et al., 2010, and our unpublished data). These findings suggest that indeed higher-ordered oligomers may be needed for full-blown IRE1α activation. They also underline an important role of the luminal domain in the activation process, probably through promotion of IRE1α dimerization and oligomerization.

### IRE1α dimers can phosphorylate other dimers in trans

The above findings suggested that phosphorylation is triggered by formation of stable oligomers or upon collisions between dimers: both events would be more frequent in cells expressing IRE1α at high concentrations. Thus, we exploited the Tet-inducible system and pushed the expression of Dim-IRE1α to higher levels. When expressed at high concentration, Dim-IRE1α became phosphorylated and was able to splice XBP1 (Fig 2B, C and Fig 3A). Considering its overexpression, the proportion of phosphorylated Fv2E-induced dimers was rather small. In absolute numbers, however, the extent of their phosphorylation was comparable to endogenous WT IRE1α. Again the level of phosphorylated IRE1α correlated with RNase activity, irrespective of the nature of the luminal domain. The same was true for the interface mutant which does not form oligomers efficiently and has no activity at moderate expression levels. At very high expression levels, also the interface mutant was phosphorylated and displayed RNase activity upon tunicamycin treatment (Fig S1C, F and Fig 3A). Thus, random collisions between dimers are important for trans-phosphorylation. Recently human IRE1α was demonstrated to form constitutive inactive homodimers at steady state (Belyy et al., 2022), which is in line with our data that formation of a dimer is not enough for self-phosphorylation.

Unlike wt IRE1α, as expected, neither Dim-IRE1α nor IM were able to form the characteristic foci upon acute stress, even if overexpressed (Li et al., 2010, and our unpublished data). Still, they were phosphorylated and acquired RNase competence upon stimulation. We can hence conclude that IRE1α molecules that are unable to form oligomers or be recruited into big clusters, can cross-phosphorylate themselves if expressed at very high levels.

In essence, the above data support the idea that a dimer can phosphorylate other dimers in trans, provided that they reach a sufficiently high local concentration. Reaching the activation threshold might entail active recruiting systems. Thus, the ability of IRE1α to form oligomers or big clusters would be key to facilitate this process. This indeed could be part of the mechanism that defends cell from spontaneous activation on IRE1α’s endonuclease.

### Phosphorylation stabilizes IRE1α dimers/oligomers in a RNase competent conformation

We reasoned that if IRE1α needs to be in higher than dimeric state to be phosphorylated, then a phosphomimetic version of Dim-IRE1α (i.e. with the three serines in the activation loop mutated to aspartic acid (Dim-S724D/S726D/S729D, in short Dim-DDD IRE1α) would be capable of splicing XBP1 upon addition of the dimerizing agent, also at much lower concentration. This was indeed the case: Dim-DDD IRE1α was able to restore XBP1 splicing, only in the presence of dimerizer and - as expected - at much lower expression levels than Dim-IRE1α (Fig 2B, C). Remarkably, the phosphorylated/phosphomimetic dimer retained full endonuclease activity. In our hands, formation of stable big oligomers was not required to splice XBP1.

### Nucleotide-binding provides an additional layer of regulation of IRE1α activity

It has been shown previously that the kinase-dead IRE1α mutant I642G can be rescued by addition of a nucleotide analog 1NM-PP1 that allosterically activates its RNase domain, mimicking adenosine nucleotide binding (Papa et al., 2003, Han et al., 2009). By the addition of 1NM-PP1, therefore, it is possible to bypass the kinase step and activate Dim-IRE1α endonuclease activity irrespectively of its phosphorylation state.

Taking advantage of this, we engineered a Dim-I642G IRE1α double mutant which allows independent control of both dimerization and RNase activation steps. As expected, Dim-I642G IRE1α displayed no phosphorylation, as revealed by phos-tag gel (Fig S3). However, the mutant was perfectly able to restore XBP1 splicing when expressed at medium expression level in the presence of the dimerizing drug and 1NM-PP1 (Fig 2C). Structural studies of the yeast Ire1p suggested that nucleotide-binding promotes oligomerization (Lee, 2008). Thus, the role of phosphorylation could be that of stabilizing the most efficient, ADP-bound, state of the protein. Sharing an identical ER luminal domain, it is unlikely that Dim-IRE1α, Dim-DDD and Dim-I642G IRE1α differ in their affinity for the dimerizer drug. Thus, if phosphorylation and subsequent formation of stable nucleotide-bound state would happen within a dimer, all 3 dimerizable mutants should be activated at similar expression levels. However, Dim-IRE1α was activated at significantly higher expression levels than Dim-DDD and Dim-I642G, implying that to be phosphorylated (and consequently ADP-bound) higher-ordered oligomers have to be formed, even if perhaps transiently. Dimerization remained a key step, regardless of phosphorylation. Indeed, in the absence of the dimerizer drug, the phosphomimetic mutant Dim-DDD IRE1 was not fully active even when expressed at high levels. Its low activity might be due to spontaneous formation of oligomers of the phosphomimetic mutant. In contrast, Dim-IRE1α requires the presence of the dimerizer drug even at very high expression levels. Taken together, these observations indicate that phosphorylation contributes to the stability of dimeric or higher-ordered structures of IRE1α, as previously suggested by structural studies (Korennykh et al., 2009) and recombinant human IRE1-kinase-endoribonuclease proteins (Le Thomas et al., 2021). The same may be true for nucleotide-binding: as shown in Fig 2C, the Dim-I642G mutant is more prone than Dim-IRE1α to display RNase activity – it is active at medium and high expression levels upon 1NM-PP1 binding even in the absence of dimerizer. These findings suggest that nucleotide-binding stabilizes IRE1α dimers and/or oligomers, which can eventually trigger RNase activity. Phosphorylation can also contribute to the stability of the nucleotide-bound state. We further confirmed it by using full-length IRE1α mutants in which kinase activity can be bypassed (I642G) and with the triple phosphomimetic mutation at S724D/S726D/S729D. Upon stress, phosphomimetic kinase bypass IRE1 (I642G-DDD) was active in the presence of 1NM-PP1 at low expression levels (Fig 3B, S1D). I642G IRE1 needed higher expression levels. This result confirmed that phosphorylation can favor formation and/or stability of higher-ordered structures of IRE1α which are more efficient in XBP1 cleavage. It was also possible that phosphomimetic I642G mutant of IRE1α had higher affinity for 1NM-PP1. Thus, dimerization-oligomerization, phosphorylation and nucleotide-binding can organize a circle of mutual support which altogether promote endonuclease activity of IRE1. Recruitment at exit sites as in case of ATF6 (Schindler and Schekman, 2009), might favor encounters between IRE1α dimers. Inhibitors would be instead BiP (Bertolotti et al., 2000) or other players that limit IRE1a congregation.

## Conclusions

The main conclusion of our study is that dimerization is not sufficient to trigger IRE1α phosphorylation. This key step in IRE1α activation must go through the formation of higher-order oligomers. According to protein–protein docking and molecular dynamics simulations, tetramers represent the most favorable configuration that IRE1α molecules can adopt (Carlesso et al., 2020). This requirement might be an important way to prevent the generation of unwanted stress signals. It seems therefore that cells need to congregate a sufficient number of IRE1α molecules in a restricted area to activate XBP1 splicing. Instead “diluting” IRE1α would lead to decreased activity of the stress sensor. Accordingly, it is of note that IRE1β suppresses endonuclease activity of IRE1α probably by forming heterocomplexes (Grey et al., 2020).

The following chain of events can be hence reconstructed for IRE1α activation (Fig 4):

1. Breaking away from BiP, IRE1α forms dimers (Bertolotti et al., 2000; Pincus et al., 2010), alternatively, it forms inactive dimers at steady state (Belyy et al., 2022)
2. Encounters between IRE1α dimers allow trans-phosphorylation
3. IRE1α phosphorylation triggers conformational changes that favor stability of higher-order oligomers (Ricci et al., 2019) and nucleotide-binding (Lee et al., 2008)
4. This in turn enhances endonuclease activity (XBP1 splicing and eventually RIDD).

**Fig 4.**
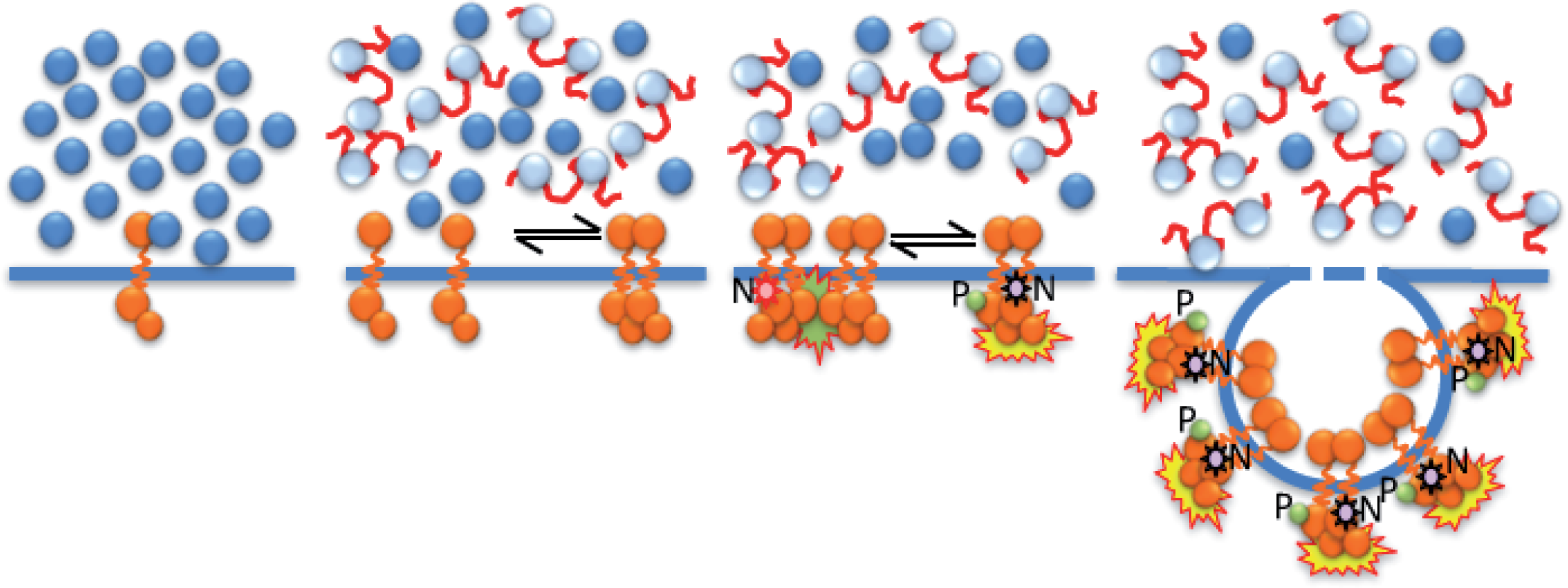
Steps of IRE1α activation. A. Schematic representation of the sequential steps that lead to full blown IRE1α activation. (A) Under basal unstressed condition, IRE1 is mostly monomeric; (B) accumulation of misfolded proteins in the ER lumen allow the formation of IRE1 dimers; (C) Collisions between dimers allow trans-phosphorylation, which enables endonuclease activity. While dimers are few or sparse, activation is mild and might be transient due to the action of phosphates, (D) Further clustering and formation of signaling foci would boost dramatically IRE1 phosphorylation by increasing local concentration of dimers, segregating them away from phosphateses, thus unleashing RIDD and proapoptotic pathways. N stands for nucleotide binding, P for phosphorylation, endonuclease activity is highlighted in yellow-red.

When ER stress is low, few dimers are formed and even fewer are able to meet and be cross-phosphorylated. At submaximal level of activation IRE1α might sustain some XBP1 splicing, but dephosphorylation could prevail. When the intensity and/or duration of stress increases, IRE1α clusters are formed that gain maximal phosphorylation and enzymatic activity. Allowing protection from phosphatase, formation of supramolecular complexes can further increase/prolong IRE1α activation, eventually driving cells into apoptosis. However, cluster formation is not necessary for XBP1 splicing.

Through the mechanism we unveil, cell responses may couple the intensity of stress and cell responses, dampening IRE1α RNase activity. IRE1α clusters would be part of the extreme measures that cells undertake when stress is overwhelming. Tilting from pro-survival to pro-apoptotic programs this mechanism would be key in stress-related diseases and a pharmacological target.

## Materials and methods

### Plasmids and IRE1α transgenes

Plasmids for lentiviral transduction of IRE1α mutant and chimeric transgenes were derived from pTETTAB in which the transgene is placed under the TetTight inducible promoter (Cohen et al. 2017).

The three IRE1α serines undergoing phosphorylation (Prischi et al., 2014) were mutated into either alanine or aspartic acid using reverse oligos `GGTACCCCtGcACGtCtcgcGAAAgcaTGCCTGCCCACTGCCAGCTT and GGTACCCCatcgCGaCGatcGAAAtcaTGaCgGCCCACTGCCAGCTTCTTGC` generating respectively S724A/S726A/S729A phospho-dead IRE1α, and S724D/S726D/S729D phosphomimetic IRE1α.

Dimerizable IRE1α was constructed by replacing the luminal domain of murine IRE1α-GFP (amino acids 1-349), or the respective IRE1α variant, with the Fv2E domain as it was done before for another ER stress sensor, Perk (Lin et al., 2009, Lu et al., 2004). The interface mutant of IRE1α, with disrupted dimerization surface, was generated by K121Y substitution according to Li et al., 2010. Finally, kinase bypass IRE1α that can be activated allosterically by NM-PP1 was obtained by I642G mutation, as described previously in (Papa et al., 2003, Han et al., 2009).

### Cell lines

Clones expressing the different IRE1α mutants at the desired levels were obtained by reconstituted HeLa-µs IRE1−KO (clone µ910-6) as described previously (Bakunts et al., 2017, Cohen et al. 2017). In brief, these cells contain a cassette for the expression of murine secretory Ig µ chains under the mifepristone inducible promoter (Sirin and

Park, 2003; Bakunts, 2017), and they no long express endogenous IRE1α due to CRISPR-mediated inactivation of the gene. Reconstitution was achieved by lentiviral transduction with the plasmids described above (Cohen et al. 2017). All IRE1α mutant genes are placed under the TetON tight promoter which allows inducible and tunable expression of the protein upon addition of Doxycycline.

Cell lines used in the paper are summarized in Supplementary file 1, including the ones specifically generated for this study: HeLa-µs IRE1α S724A/S726A/S729A (AAA), HeLa-µs IRE1α K121Y (K121Y), HeLa-µs Fv2E-IRE1α (Dim-IRE1), HeLa-µs Fv2E-IRE1α S724D/S726D/S729D (Dim-DDD) and HeLa-µs Fv2E-IRE1α I642G (Dim-I642G), HeLa-µs IRE1α N906A (RNase-dead), HeLa-µs IRE1α I642G (Kinase bypass, I642G), HeLa-µs IRE1α I642G S724D/S726D/S729D (I642G-DDD).

All cell lines in this study were ultimately derived from HeLa S3 cells, of which the genotype was confirmed by PCR single locus technology (Bakunts et al., 2017).

### Reagents and treatments

The expression of µs under the MifOn promoter was triggered by treatment with 0.5 nM Mifepristone (Bakunts et al. 2017). The expression of IRE1α variants was generally induced with Doxycycline for 48-72 hours before the experiment. The dose of Doxycycline to use was determined empirically for each individual cell line and adjusted in order to achieve close-to-endogenous (low), medium or high levels of expression, as determined by comparative western blots.

For allosteric activation of I642G IRE1α, 1NM-PP1 (MedchemExpress) was used at the concentration of 7 µM. Dimerization of Fv2E domains was induced by treatment with AP20187 (MedchemExpress), at a concentration of 10 nM, unless stated otherwise. Kifunensin (Sigma-Merck) was used at the concentration of 30 µM.

Rabbit-anti-human IRE1α was from Cell signaling (3294) and rabbit-α-human phosphoIRE1α (S724) was from Abcam (ab48187).

### Analysis of IRE1α phosphorylation and activity

To detect IRE1α phosphorylation we used the method previously described in (Yang et al., 2010). Cells were lysed in 50 mM Tris-HCl, 150 mM NaCl, 60mM octylglucoside, pH 7.4 containing phosphatase inhibitors cocktails 2 and 3 (Sigma). Proteins were separated by SDS-PAGE on a 5% Polyacrylamide gel containing 20 μM Phos-tag (NARD, Wako-chemicals), performed at 15 mA for 2-2.5 h. The gel was transferred to nitrocellulose (Bio-Rad) and decorated with anti-IRE1α antibodies.

IRE1α RNase activity was determined by XBP1 splicing assay, as described previously (Bakunts et al. 2017).

## Abbreviations

UPR: unfolded protein response
ER: endoplasmic reticulum
ERAD: ER-associated degradation
IRE1: Inositol-requiring enzyme type 1

## Acknowledgements

We thank David Ron for help with reagents, Tiziana Anelli for suggestions and Giuliano Martino, Laura Tadè and Silvia Russo Krauss for help with experiments.

This work was supported in part via grants from AIRC (IG 2019 - ID. 23285) and MUR (PRIN 2017XA5J5N).

## Legends to Supplement Figures

**Fig S1.**
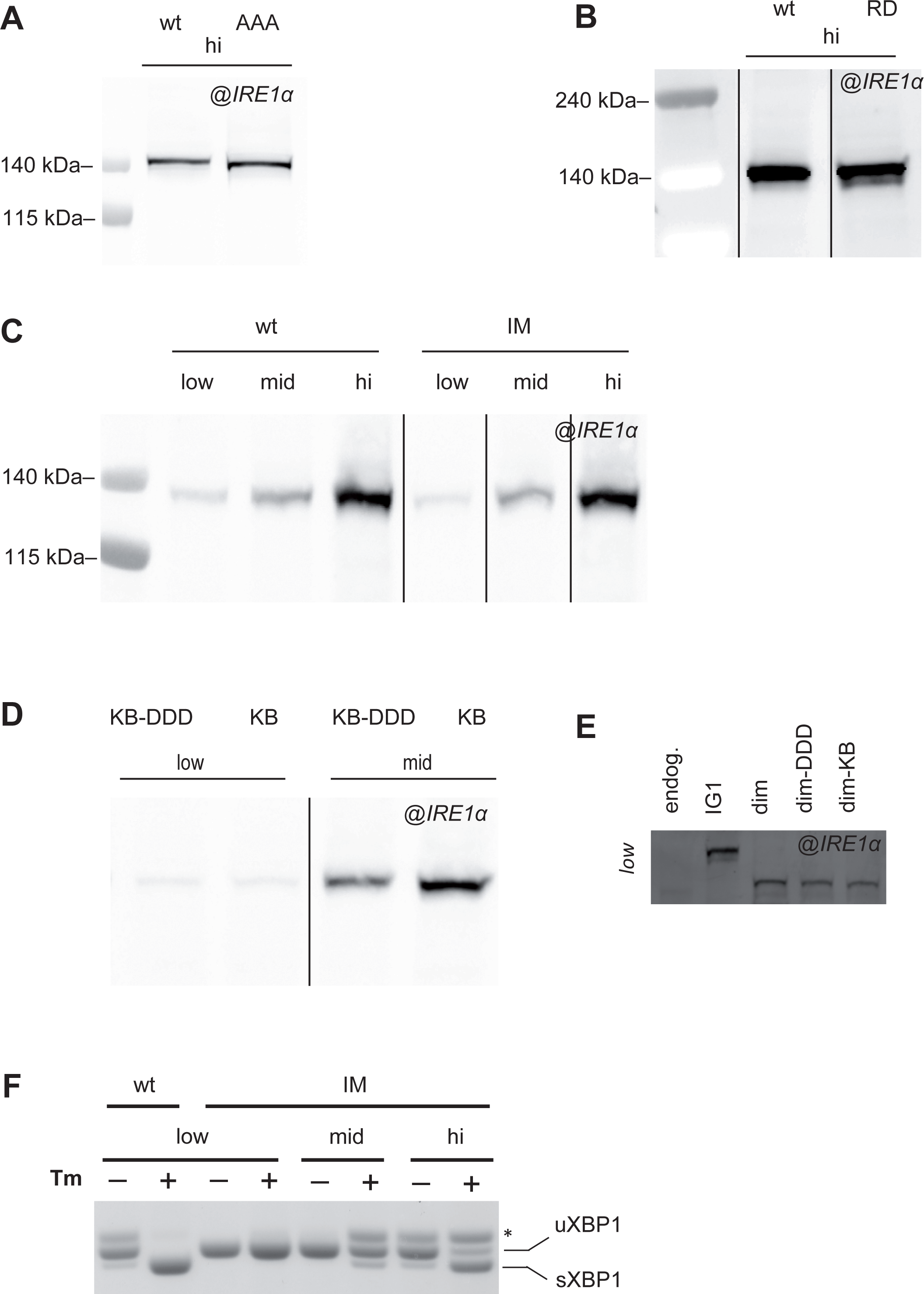
A-D. HeLa clones expressing IRE1α mutants under the TetON inducible promoter were treated with different concentrations of Doxycycline to adjust the expression level of the transgenes (Vitale M et al. 2019). Aliquots from the cell lysates were resolved electrophoretically and the blots decorated with anti-IRE1α. (A) Phospho-incompetent AAA mutant, (B) N906A RNAse dead mutant, (C) K121Y interface mutant; (D) I642G-DDD, kinase bypass phosphomimetic mutant. E. Comparison of the expression of IRE1α mutants at low level (higher exposure of “low” panel of Fig 2B). F. Endonuclease activity of IRE1α interface mutant at different expression levels was assessed by XBP1 splicing assay. Clearly, the mutant is active only at high expression levels.

**Fig S2.**
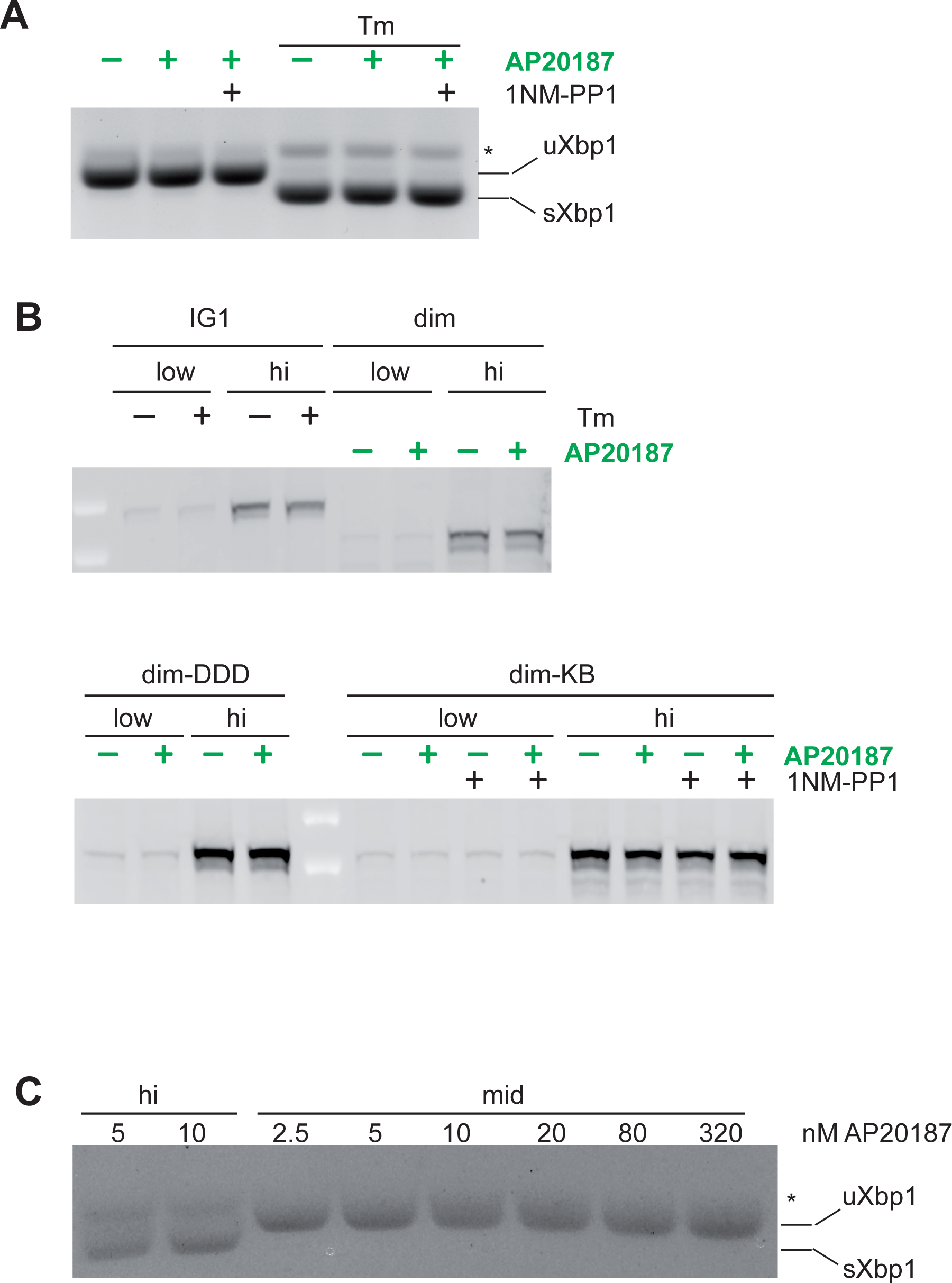
A. Neither the dimerizer drug (AP20187) nor 1NM-PP1 affect XBP1 the activity of endogenous IRE1α in HeLa-µ_s_ cells. Cells were treated for 4 h with or without the drugs as indicated. B. Neither Tm, nor AP20187, nor 1NM-PP1 impact the expression levels of wt (IG1) and dimerizable IRE1 (dim) (upper panel) neither those of dime-DDD and dim-I642G (lower panel) mutants. C. Increasing concentrations of AP20187 do not change RNase activity of Dim-IRE1. 10nM of dimerizer drug AP20187 was used for the experiments, as described in the text.

**Fig S3.**
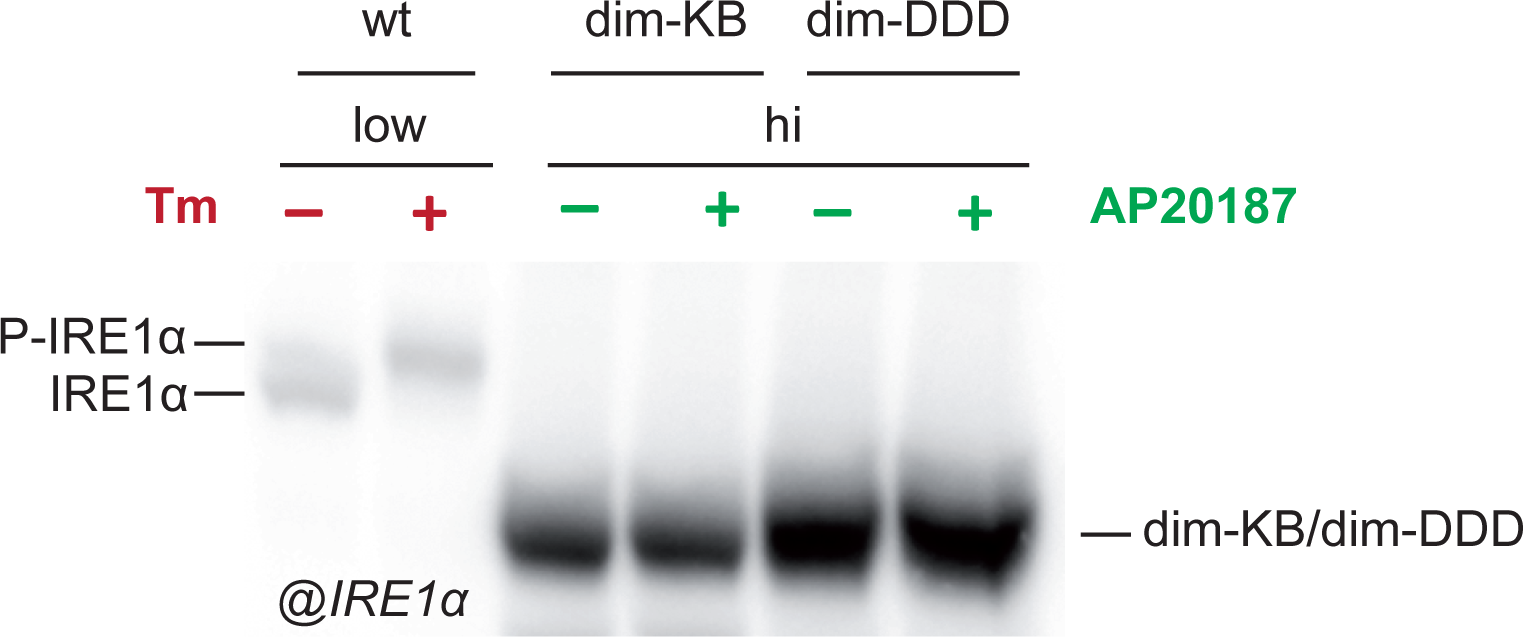
As expected, neither the phosphomimetic Dim-DDD (in which aspartic acid replaces the key 3 serines) nor Dim-I642G (due to the inability to bind ATP and, hence, phosphotransfer) can be phosphorylated even if highly expressed. Immunoblots of IRE1α using the Phos-tag gels with anti-IRE1 and anti-IRE1 (phospho S724) antibodies.

## References

Anelli T, Sitia R (2008) Protein quality control in the early secretory pathway. EMBO J 27.315–27

Aragón T, van Anken E, Pincus D, Serafimova IM, Korennykh AV, Rubio CA, Walter P (2009) Messenger RNA targeting to endoplasmic reticulum stress signalling sites. Nature 457: 736–740

Bakunts A, Orsi A, Vitale M, Cattaneo A, Lari F, Tadè L, Sitia R, Raimondi A, Bachi A, van Anken E (2017) Ratiometric sensing of BiP-client versus BiP levels by the unfolded protein response determines its signaling amplitude. eLife 6: e27518

Belyy V, Zuazo-Gaztelu I, Alamban A, Ashkenazi A, Walter P (2022) Endoplasmic reticulum stress activates human IRE1α through reversible assembly of inactive dimers into small oligomers. Elife 11: e74342

Bertolotti A, Zhang Y, Hendershot LM, Harding HP, Ron D (2000) Dynamic interaction of BiP and ER stress transducers in the unfolded-protein response. Nat. Cell Biol 2: 326–332

Carlesso A, Hörberg J, Reymer A, Eriksson LA (2020) New insights on human IRE1 tetramer structures based on molecular modeling. Sci Rep 10: 17490

Cohen N, Breker M, Bakunts A, Pesek K, Chas A, Argemi J, Orsi A, Gal L, Chuartzman S, Wigelman Y, Jonas F, Walter P, Ernst R, Aragon T, van Anken E, Schuldiner M (2017) Iron affects Ire1 clustering propensity and the amplitude of endoplasmic reticulum stress signaling. JCS 130: 3222–3233

Ellgaard, L, Helenius, A (2003) Quality control in the endoplasmic reticulum. Nat Rev Mol Cell Biol 4:181–91

Grey MJ, Cloots E, Simpson MS, LeDuc N, Serebrenik YV, De Luca H, De Sutter D, Luong P, Thiagarajah JR, Paton AW, Paton JC, Seeliger MA, Eyckerman S, Janssens S, Lencer WI (2020) IRE1β negatively regulates IRE1α signaling in response to endoplasmic reticulum stress. J Cell Biol 219: e201904048

Han D, Lerner AG, Vande WL, Upton JP, Xu W, Hagen A, Backes BJ, Oakes SA, Papa FR (2009) IRE1alpha kinase activation modes control alternate endoribonuclease outputs to determine divergent cell fates. Cell 138: 562–575

Hollien J, Lin JH, Li H, Stevens N, Walter P, Weissman JS (2009) Regulated Ire1-dependent decay of messenger RNAs in mammalian cells. JCB 186: 323–331

Korennykh AV, Egea PF, Korostelev AA, Finer-Moore J, Zhang C, Shokat KM, Stroud RM, Walter P (2009) The unfolded protein response signals through high-order assembly of Ire1. Nature 457: 687–93

Korennykh AV, Korostelev AA, Egea PF, Finer-Moore J, Stroud RM, Zhang C, Shokat KM, Walter P (2011) Structural and functional basis for RNA cleavage by Ire1. BMC Biol 9: 47

Korennykh A, Walter P (2012) Structural basis of the unfolded protein response. Annu Rev Cell Dev Biol 28: 251–77

Krshnan L, van de Weijer ML, Carvalho P (2022) Endoplasmic Reticulum-Associated Protein Degradation. Cold Spring Harb Perspect Biol 14: a041247

Le Thomas A, Ferri E, Marsters S, Harnoss JM, Lawrence DA, Zuazo-Gaztelu I, Modrusan Z, Chan S, Solon M, Chalouni C, Li W, Koeppen H, Rudolph J, Wang W, Wu TD, Walter P, Ashkenazi A (2021) Decoding non-canonical mRNA decay by the endoplasmic-reticulum stress sensor IRE1α. Nature Communications 12: 1–15

Lee KP, Dey M, Neculai D, Cao C, Dever TE, Sicheri F (2008) Structure of the Dual Enzyme Ire1 Reveals the Basis for Catalysis and Regulation in Nonconventional RNA Splicing. Cell 132: 89–100

Li H, Korennykh AV, Behrman SL, Walter P (2010) Mammalian endoplasmic reticulum stress sensor IRE1 signals by dynamic clustering. PNAS 107: 16113–16118

Lin JH, Li H, Zhang Y, Ron D, Walter P (2009) Divergent Effects of PERK and IRE1 Signaling on Cell Viability. PLoS One 4: e4170

Lu PD, Jousse C, Marciniak SJ, Zhang Y, Novoa I, Scheuner D, Kaufman RJ, Ron D, Harding HP (2004) Cytoprotection by pre-emptive conditional phosphorylation of translation initiation factor 2. EMBO J 23: 169–179

Morita S, Villalta SA, Feldman HC, Register AC, Rosenthal W, Hoffmann-Petersen IT, Mehdizadeh M, Ghosh R, Wang L, Colon-Negron K, Meza-Acevedo R, Backes BJ, Maly DJ, Bluestone JA, Papa FR (2017) Targeting ABL-IRE1α signaling spares ER-stressed pancreatic β cells to reverse autoimmune diabetes. Cell Metab 25: 883–897

Papa FR, Zhang C, Shokat K, Waiter P (2003) Bypassing a Kinase Activity with an ATP-Competitive Drug. Science 302: 1533–1537

Pincus D, Chevalier MW, Aragon T, van Anken E, Vidal SE, El-Samad H, Walter P (2010) BiP binding to the ER-stress sensor Ire1 tunes the homeostatic behavior of the unfolded protein response. PLoS Biol 8: e1000415

Prischi F, Nowak PR, Carrara M, Ali MMU (2014) Phosphoregulation of Ire1 RNase splicing activity. Nature Communications 5: 3554

Raymundo DP, Doultsinos D, Guillory X, Carlesso A, Eriksson LA (2020) Pharmacological Targeting of IRE1 in Cancer. Trends Cancer 6: 1018–1030

Ricci D, Marrocco I, Blumenthal D, Dibos M, Eletto D, Vargas J, Boyle S, Iwamoto Y, Chomistek S, Paton JC, Paton AW, Argon Y (2019) Clustering of IRE1α depends on sensing ER stress but not on its RNase activity. FASEB J 33: 9811–9827

Schindler AJ and Schekman R (2009) In vitro reconstitution of ER-stress induced ATF6 transport in COPII vesicles. PNAS 106: 17775–17780

Shang J, Lehrman MA (2004) Discordance of UPR signaling by ATF6 and Ire1p-XBP1 with levels of target transcripts. Biochemical and Biophysical Research Communications 317: 390–396

Sirin O, Park F (2003) Regulating gene expression using self-inactivating lentiviral vectors containing the mifepristone-inducible system. Gene 323:67–77

Sitia R, Braakman I (2003) Quality control in the endoplasmic reticulum protein factory. Nature 426: 891–894

Sundaram A, Appathurai S, Plumb R, Mariappan M (2018) Dynamic changes in complexes of IRE1α, PERK, and ATF6α during endoplasmic reticulum stress. Mol Biol Cell 29: 1376–1388

Vitale M, Bakunts A, Orsi A, Lari F, Tadè L, Danieli A, Rato C, Valetti C, Sitia R, Raimondi A, Christianson JC, van Anken E (2019) Inadequate BiP availability defines endoplasmic reticulum stress. eLife 8: e41168

van Anken E, Pincus D, Coyle S, Aragón T, Osman C, Lari F, Puerta SG, Korennykh AV, Walter P (2014) Specificity in endoplasmic reticulum-stress signaling in yeast entails a step-wise engagement of HAC1 mRNA to clusters of the stress sensor Ire1. eLife 3: e05031

Walter P, Ron D (2011) The unfolded protein response: from stress pathway to homeostatic regulation. Science 334: 1081–6

